# Sub-Concussive Head Impacts From Heading Footballs Do Not Acutely Alter Brain Excitability As Compared to a Control Group

**DOI:** 10.1101/2023.05.31.543027

**Authors:** R. Hamel, B. M. Waltzing, T. Massey, J. Blenkinsop, L. McConnell, K. Osborne, K. Sesay, F. Stoneman, A. Carter, H. Maaroufi, N. Jenkinson

**Affiliations:** School of Sports, Exercise, and Rehabilitation Sciences, University of Birmingham, United Kingdom; Institute of Neurosciences, UC Louvain, Belgium Avenue Mounier 54, 1200, Bruxelles, Belgium

**Keywords:** Corticomotor inhibition, cortical silent period, transcranial magnetic stimulation, sub-concussive head impacts, brain excitability

## Abstract

**Background:** Repeated sub-concussive head impacts are a growing brain health concern, but the possible mechanisms of trauma and plausible biomarkers remain elusive. One impediment is the lack of an experimental model to study the effects of sub-concussive head impacts on the brain.

**Objectives:** This work’s objective was to provide an experimental model to study the acute effects of sub-concussive head impacts on the brain. To do so, this study aimed to replicate previous work from Di Virgilio et al. (2016) showing that head impacts from heading footballs acutely alter brain excitability by increasing corticomotor inhibition.

**Methods:** Scores from the Rivermead Post-Concussion Questionnaire and measurements of cortical silent period (CSP) duration – obtained using transcranial magnetic stimulation to assess corticomotor inhibition in the central nervous system – were taken before and after participants performed 20 football headings (Headings; n = 30) or control (Control; n = 30).

**Results:** The results revealed increased headaches and dizziness symptoms in the Headings as compared to the Control group, revealing the qualitative experience of head impacts. The results then revealed that CSP duration similarly lengthened in both the Headings and Control groups, suggesting that head impacts did not cause the increased corticomotor inhibition.

**Conclusions:** The results show that head impacts from football headings did not acutely alter corticomotor inhibition as compared to a control group that did not experience head impacts, suggesting that excitability changes do not reflect acute sub-concussive brain injuries. Nonetheless, this work suggests that football headings can be used as an experimental model to study the effects of sub-concussive head impacts on brain health. Future work could use the present procedures to investigate additional biomarkers of brain injury.

## Introduction

Repeated sub-concussive head impacts are increasingly recognised to have detrimental effects on brain health (Ntikas et al., 2022). For instance, neuroimaging studies have shown that repeated sub-concussive head impacts negatively alter white matter tract integrity, reduce cortical thickness, and impair cognitive functions (Koerte et al., 2015; Lipton et al., 2013). Moreover, a chronic history of sub-concussive head impacts – such as those experienced by football players when headings footballs – increases the risk of developing neurodegenerative diseases (Russell et al., 2021; Ueda et al., 2023). Given that the socioeconomic burden of neurodegenerative diseases is increasing worldwide (Wong, 2020), and that head impacts are modifiable risk factors (Livingston et al., 2020), there is a pressing need to characterize the full effects of sub-concussive head impacts on brain health using randomised controlled experimental settings (Batty & Kaprio, 2022). The overarching objective of this work was to provide an experimental model to study the acute effects of sub-concussive head impacts on the brain. Similar to work on concussions (Feddermann-Demont et al., 2020), such systematic investigations could ultimately lead to recommendations to attenuate (or manage) the detrimental effects of head impacts on brain health.

One promising model is to use transcranial magnetic stimulation (TMS) to non-invasively assess changes in brain excitability in response to performing football headings (Di Virgilio et al., 2016; Ntikas et al., 2022). Specifically, Di Virgilio et al. (2016) used TMS to measure changes in the cortical silent period (CSP) duration – defined as the suppression of voluntary electromyography activity, which linearly reflects the extent of GABAergic corticomotor inhibition in the corticospinal tract (Ziemann et al., 2015) – before and after participants performed 20 football headings over 10min. The authors found that CSP duration lengthened immediately after performing the headings, suggesting that sub-concussive head impacts acutely altered brain excitability by increasing corticomotor inhibition. Although these results were crucially not compared to a control group that did not experience head impacts, this approach remains a promising model because it allows to experimentally induce sub-concussive head impacts in a controlled environment. Furthermore, changes in CSP duration constitute a promising biomarker, as measuring CSP is non-invasive, can be easily and quickly measured, and its utility to evaluate changes in brain excitability is supported by work on both concussive (Scott et al., 2020) and sub-concussive head impacts (Ntikas et al., 2022). Overall, this evidence suggests that measuring changes in CSP duration before and after performing football headings constitutes a promising experimental model.

Therefore, the objective of this study was to replicate the results from Di Virgilio et al. (2016) using a randomised controlled trial – by adding an appropriate control group – to ascertain that changes in CSP duration reflect acute brain injuries. This work also sought to confirm that football headings can be used as an experimental model to study the acute effects of sub-concussive head impacts on brain functions. Here, scores from Rivermead Post-Concussion Questionnaire and changes in CSP duration were assessed before and immediately after groups of participants performed (Headings; n = 30), or not (Control; n = 30), 20 football headings. It was hypothesized that CSP would acutely lengthen in response to the head impacts (as in Di Virgilio et al., 2016) *and* as compared to the control group.

## METHODS

### Participants

Sixty physically and neurologically healthy young adults took part in this study (mean ± 95% confidence intervals (CIs) = 20.9 ± 0.7 years old; hereafter, all descriptive statistics represent the mean ± 95% CIs). All participants reported having no history of concussion in the previous 5 years, suggesting preserved integrity of neurophysiological activity within the motor cortex (Tremblay et al., 2014). To enhance the sample’s ecological validity, participants with or without expertise playing in football were recruited. It was reasoned that the effects of sub-concussive head impacts on the brain’s excitability should be apparent regardless of players’ expertise (see Di Virgilio et al., 2022), so football headings can be employed as a valid experimental model.

The participants were randomly allocated to a group that experienced sub-concussive head impacts (Headings; n = 30; 5 females; 20.9 ± 1.1 years) and a group that did not (Control; n = 30; 5 females; 20.8 ± 0.9 years). Participants were screened for TMS contraindications (Rossi et al., 2009), the major contraindications being the presence of metal particles/objects in the region of the head and the presence of a personal history of epileptic seizures or convulsions. The study was approved by the local research ethics committee of the University of Birmingham (project # ERN-182077AP10). All participants provided written informed consent before their participation.

The number of participants was based on a sample size analysis using G*Power (v.3.1.9.4). Namely, the smallest effect size of interest for the context of this study was a Cohen’s d of 0.8 (large effect size). Assuming the use of an independent t-test (to evaluate differences between Headings and Control), 80% statistical power, and a significance threshold of 0.05, the results of the analysis revealed that two groups of 26 participants are required. To ensure a statistical power greater than 80%, two groups of 30 participants were thus recruited. Furthermore, a post-hoc achieved power analysis using G*Power was conducted to determine if the statistical power of this study would be enough to replicate the acute lengthening in CSP duration from Di Virgilio et al. (2016), which reported a Cohen’s dz of 1.125 (above-large effect size). Assuming this effect size, a significance threshold of 0.05, and a sample size of 30 participants, the achieved power in the present study was 100%. Overall, these results suggest that two groups of 30 participants are sufficient to detect meaningful differences between the Headings and Control groups and replicate the results from Di Virgilio et al. (2016).

### Study protocol

An overview of the study protocol is provided in Figure 1. The Rivermead Post-Concussion Questionnaire, the CSP (Figure 1A) and the exerted force during a voluntary contraction were measured before (Pre Measures) and after (Post Measures; Figure 1C) participants either headed the ball or used their foot to control the ball (Figure 1B). In both groups, participants experienced 20 throws, at a rate of 1 throw every 30 sec (as in Di Virgilio et al., 2016). Trials where participants missed the ball were redone to ensure that a total of 20 contacts were experienced with the ball. Participants of the Headings group were instructed to head the football to redirect it perpendicularly in the direction of their right-hand side (rotational headers; as in Di Virgilio et al., 2016). Participants of the Control group were instructed to use their dominant lower limb (preferably the foot) to control the ball. This method of control was used to prevent head impacts and match the level of exercise performed between the two groups. All parts of the procedures involving footballs were performed outdoors.

**Figure 1.**
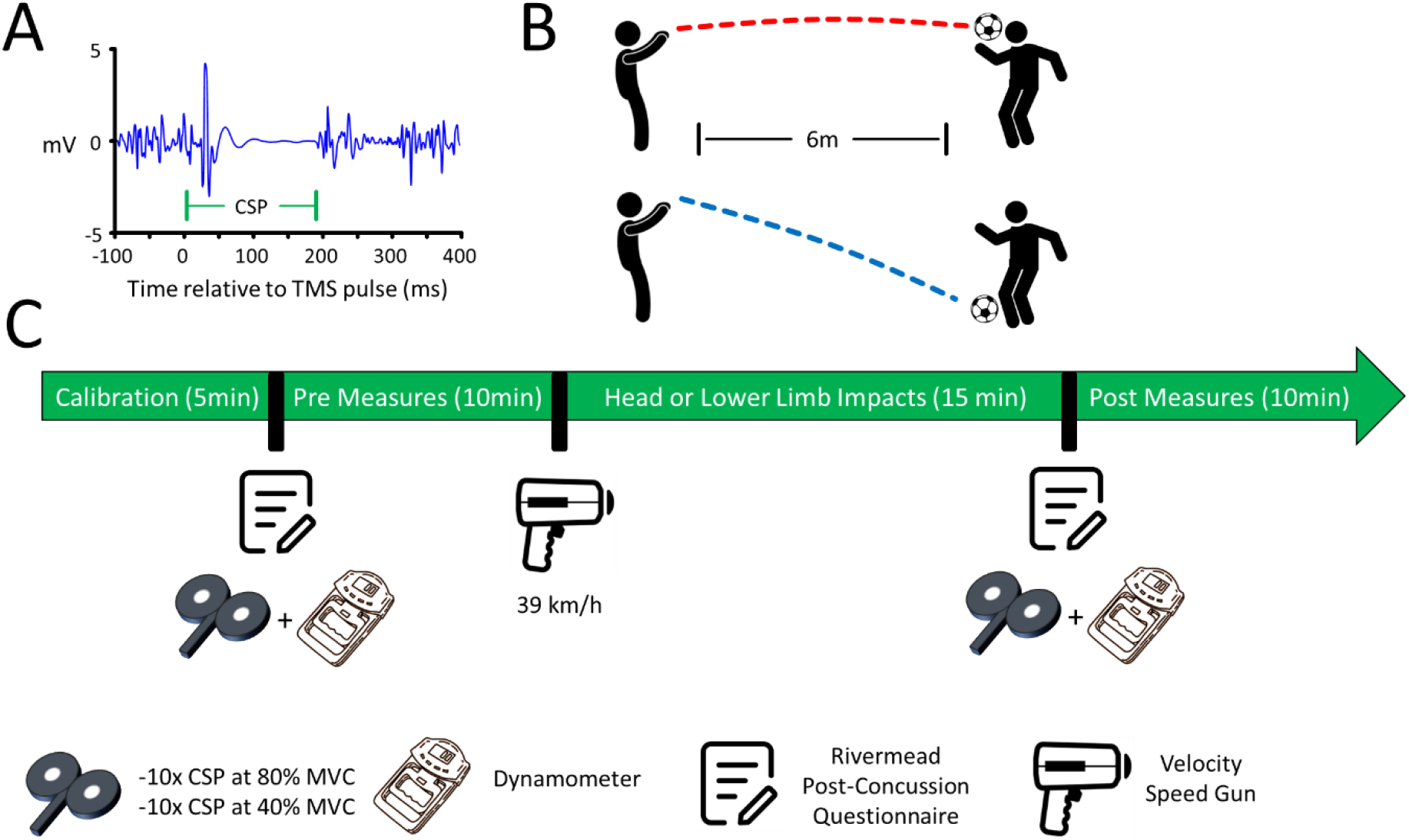
Overview of the protocol. **(A)** *Visual depiction of a cortical silent period (CSP)*. A silent period in the surface electromyography (EMG) follows the delivery of a TMS pulse when participants are voluntarily contracting hand muscles. The CSP duration is thought to positively covary with the extent of GABAergic inhibition in the corticospinal tract (Ziemann et al., 2015). **(B)** *Visual depiction of the Headings and Control groups*. An experienced experimenter manually threw the ball from a 6m distance at a set speed of 39 km/h (measured using a velocity speed gun on every throw). The ball was thrown a total of 20 times (once every 30 sec). Participants either headed the ball (Headings; upper panel) or controlled the ball using their dominant lower limb (Control; lower panel), therefore preventing any head impacts. **(C)** *Timeline of the protocol*. The experiment began with the calibration of the surface EMG, TMS and dynamometer. Then, participants filled out the Rivermead Questionnaire and a total of 20 valid CSP trials were collected (10 trials at 80% of the maximum voluntary contraction (MVC) and 10 trials at 40% MVC). Subsequently, participants performed the headings or lower limb ball contacts. Finally, the Rivermead Questionnaire was filled and a total of 20 valid CSP trials were collected a second time (10 trials at 80% MVC and 10 trials at 40% MVC). The measures were collected in this specific order for every participant.

The ball consisted of a standard football (400g, 70 cm circumference; 8 psi) and was manually thrown by an experienced (> 10 years) football player from a 6m distance (Di Virgilio et al., 2016) towards the head (Headings) or dominant lower limbs (aiming at the foot; Control). The rationale for manually throwing the ball was to best approximate practice and game conditions, as an effort to enhance ecological validity. The ball was thrown at an average speed of 39km/h (Di Virgilio et al., 2016), which was assessed upon each throw with a hand-held Velocity Speed Gun (Bushnell, model 101911). Ball speed for the Headings and Control groups was 39.15 ± 0.16 km/h and 39.09 ±0.14 km/h, respectively, confirming that the 39km/h target was successfully achieved. Globally, 40min were required to complete the protocol (including the questionnaires and TMS procedures; Figure 1C).

### Questionnaire

To assess sub-concussive head impact symptoms, the Rivermead Post-Concussion Questionnaire was used (Potter et al., 2006). The Rivermead Questionnaire consists of 16 items (see Table 1 for a list), each evaluated using the following scores: 0 (not experienced at all), 1 (no more of a problem), 2 (a mild problem), 3 (a moderate problem), and 4 (a severe problem). Conceivably, some items are unspecific in evaluating symptoms of acute sub-concussive head impacts (e.g., feeling frustrated, sleep disturbance), but other items arguably appear well-suited to do so (e.g., headaches, feeling of dizziness). In the absence of a validated questionnaire specifically developed for that purpose, it was reasoned that the Rivermead Questionnaire would be sensitive enough to evaluate some of the symptoms of sub-concussive head impacts. The results provide post-hoc support for this contention.

**Table 1.** Results of the Rivermead Questionnaire – Mean (95% CIs)

|  | Headings |  | Control |  |
| --- | --- | --- | --- | --- |
|  | Pre | Post | Pre | Post |
| Headaches | 0.033<br>(0.065) | 0.767<br>(0.292) | 0.100<br>(0.109) | 0.033<br>(0.065) |
| Feeling of dizziness | 0.000<br>(0.000) | 0.300<br>(0.191) | 0.000<br>(0.000) | 0.033<br>(0.065) |
| Nausea/vomiting | 0.000<br>(0.000) | 0.100<br>(0.144) | 0.000<br>(0.000) | 0.000<br>(0.000) |
| Noise sensitivity | 0.000<br>(0.000) | 0.033<br>(0.065) | 0.033<br>(0.065) | 0.000<br>(0.000) |
| Sleep disturbance | 0.167<br>(0.165) | 0.133<br>(0.155) | 0.133<br>(0.155) | 0.100<br>(0.144) |
| Fatigue, tiring more easily | 0.167<br>(0.165) | 0.200<br>(0.146) | 0.133<br>(0.155) | 0.100<br>(0.144) |
| Irritable, easily angered | 0.033<br>(0.065) | 0.033<br>(0.065) | 0.000<br>(0.000) | 0.000<br>(0.000) |
| Depressed or tearful | 0.067<br>(0.091) | 0.033<br>(0.065) | 0.100<br>(0.196) | 0.067<br>(0.131) |
| Frustrated or impatient | 0.033<br>(0.065) | 0.033<br>(0.065) | 0.100<br>(0.109) | 0.033<br>(0.065) |
| Forgetfulness, poor memory | 0.067<br>(0.091) | 0.067<br>(0.091) | 0.033<br>(0.065) | 0.033<br>(0.065) |
| Poor concentration | 0.100<br>(0.109) | 0.233<br>(0.180) | 0.200<br>(0.173) | 0.133<br>(0.155) |
| Taking longer to think | 0.067<br>(0.091) | 0.200<br>(0.173) | 0.033<br>(0.065) | 0.000<br>(0.000) |
| Blurred vision | 0.000<br>(0.000) | 0.033<br>(0.065) | 0.000<br>(0.000) | 0.000<br>(0.000) |
| Light sensitivity | 0.033<br>(0.065) | 0.033<br>(0.065) | 0.000<br>(0.000) | 0.000<br>(0.000) |
| Double vision | 0.000<br>(0.000) | 0.000<br>(0.000) | 0.000<br>(0.000) | 0.000<br>(0.000) |
| Restlessness | 0.000<br>(0.000) | 0.067<br>(0.131) | 0.100<br>(0.109) | 0.100<br>(0.109) |

### TMS, EMG, and Dynamometer

The following procedures were performed indoors, in a temperature-controlled environment. TMS was delivered using a 70mm figure-of-eight D-Alpha flat coil connected to a Magstim 200^2^ stimulator (Magstim, Whitland, UK). TMS pulses were delivered to the cortical representation of the first dorsal interosseus (FDI) muscle of the left M1 (McNabb et al., 2020). The surface electromyography (EMG) data of the FDI muscle of the right hand were recorded using bipolar electrodes connected to a Delsys ® Bagnoli system, itself connected to a Power 1402 data acquisition interface (Cambridge Electronic Design ®). The data were digitised at 10,000 Hz for 500ms epochs (100ms pre-trigger time; bandpass filtered between 20 and 450 Hz) and recorded using Signal (v6.05; Cambridge Electronic Design ®). The reference electrode was located on the proximal portion of the right ulnar bone.

Participants wore a rubber swimming cap, which was used to mark the location of the hotspot (Hamel et al., 2020). Vertical lines were additionally drawn at the junction of participants’ skin and swimming cap, to ensure that the swimming cap did not move when performing the headings. The hotspot location, defined as the cortical location where motor-evoked potentials (MEPs) in the FDI were reliably induced, was found by delivering suprathreshold pulses. Then, the stimulator intensity was adjusted to obtain MEPs of ±1mV at rest. For all participants (n = 60), the stimulator intensity was set at 55 ± 2%. This was defined as the test stimulus intensity and was kept constant for the experiment’s duration for a given participant.

A total of 10 valid CSP trials per level of maximum voluntary contraction (MVC; see below) for both Pre and Post Measures were recorded. This number of trials was chosen because measuring more than 10 CSP trials does not enhance the reliability of CSP duration estimation (Garvey et al., 2001). To induce CSP, participants squeezed a hand dynamometer (Kuptone ®) at 80% and 40% of their MVC (defined as the greatest force generated out of three trials). Although arguably non-specific to the FDI muscle, the use of a dynamometer was to provide an effective, inexpensive, and accessible means to induce voluntary contractions and measure MVC of the hand muscles for on-the-field and experimental settings. Participants were instructed to reach and maintain a contraction at their individualised 80% and 40% MVC targets for ∼3sec, allowing sufficient time for the experimenter to deliver a single TMS pulse upon reaching those targets. The peak generated force (in kg) was assessed for each CSP trial and kept for analysis. The results revealed that all participants (n = 60) exerted peak forces that represented 80 ± 0.1% and 43 ± 0.1% of their MVC. A target of 100% MVC was not used (unlike Di Virgilio et al., 2016), as it generated important fatigue and limited the number of CSP trials that could be recorded. CSP trials were collected at 80% and 40% MVC to determine if similar CSP lengthening could be observed at different MVC (as supported by Kojima et al., 2013). The case being, it would suggest that CSP duration lengthening in response to head or lower limb impacts (Figure 1) could be observed using low MVC, making it convenient to collect multiple trials to reliably evaluate the effects of sub-concussive head impacts on CSP duration.

### Dependent variables

The main dependent variables were CSP duration and the Rivermead Questionnaire scores. Using a custom-designed algorithm in MatLab, CSP duration was measured as the time difference (in ms) between the delivery of the TMS pulse and the return of the EMG of voluntary muscle activity (CSP offset). Specifically, the CSP offset was determined as the moment when the SD of a 2.5ms sliding window exceeded 50% of the SD of the EMG background activity, calculated over the 100ms that preceded TMS pulse delivery, for at least 5ms (similar to Goodall et al., 2010; Hamel et al., 2022). The EMG data were not rectified. To normalise the CSP duration, the CSP data (in ms) from the Post Measures were divided by the CSP data (in ms) from the Pre Measures. This was done separately for each level of Contraction Levels (80%, 40% MVC). The Rivermead Questionnaire scores (0 to 4) were averaged across groups for each item (see Table 1).

### Statistical analysis

To analyse the results, mixed ANOVAs were used. The within-subject factors were Measures (Pre, Post) and Contraction Levels (80% MVC, 40% MVC). The between-subject factor was Groups (Headings, Control). If the data violated the assumptions of sphericity (*p* < 0.05, Mauchly test), the Greenhouse-Geiser correction was applied. If data deviated from normality (p < 0.05; Shapiro-Wilk test) upon pairwise comparisons, non-parametric pairwise comparisons were conducted (Wilcoxon rank test rather than dependent t-test for within-subject comparisons; U Mann-Whitney test rather than independent t-test for between-subject comparisons). The Benjamini-Hochberg (1995) correction was used to control for inflated type 1 error upon multiple comparisons. The statistical significance threshold was set at 0.05. All descriptive statistics reported in this work represent the mean ± 95% CIs. The open-access software JAMOVI was used to conduct the statistical analyses.

## Results

### Increased headaches and dizziness symptoms after football heading

The data from the 16-item Rivermead Questionnaire were analysed using 2 Measures (Pre, Post) * 2 Groups (Headings, Control) mixed ANOVAs. See Table 1 for a complete report of the group questionnaire data. Overall, the results below show that heading footballs significantly increased headaches and dizziness symptoms as compared to the control group, providing post-hoc support to using the Rivermead Questionnaire to assess some of the symptoms of sub-concussive head impacts.

For headaches symptoms (Figure 2A), the results revealed a Measures * Groups interaction (F_(1,58)_ = 23.460, *p* < 0.0001, 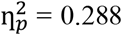). Breakdown of the Measures * Groups interaction revealed that headaches symptoms increased from Pre to Post for the Headings group (*p* = 0.0008, Cohen’s dz = 0.845), but not for the Control group (*p* = 0.3458, Cohen’s dz = 0.263). The Headings and Control groups did not differ at Pre (*p* = 0.3128, Cohen’s dz = 0.265), but the Headings showed greater headaches symptoms than the Control group at Post (*p* < 0.0001, Cohen’s d = 1.239). This analysis indicates that participants experienced headaches symptoms when heading footballs but not in the control group.

**Figure 2.**
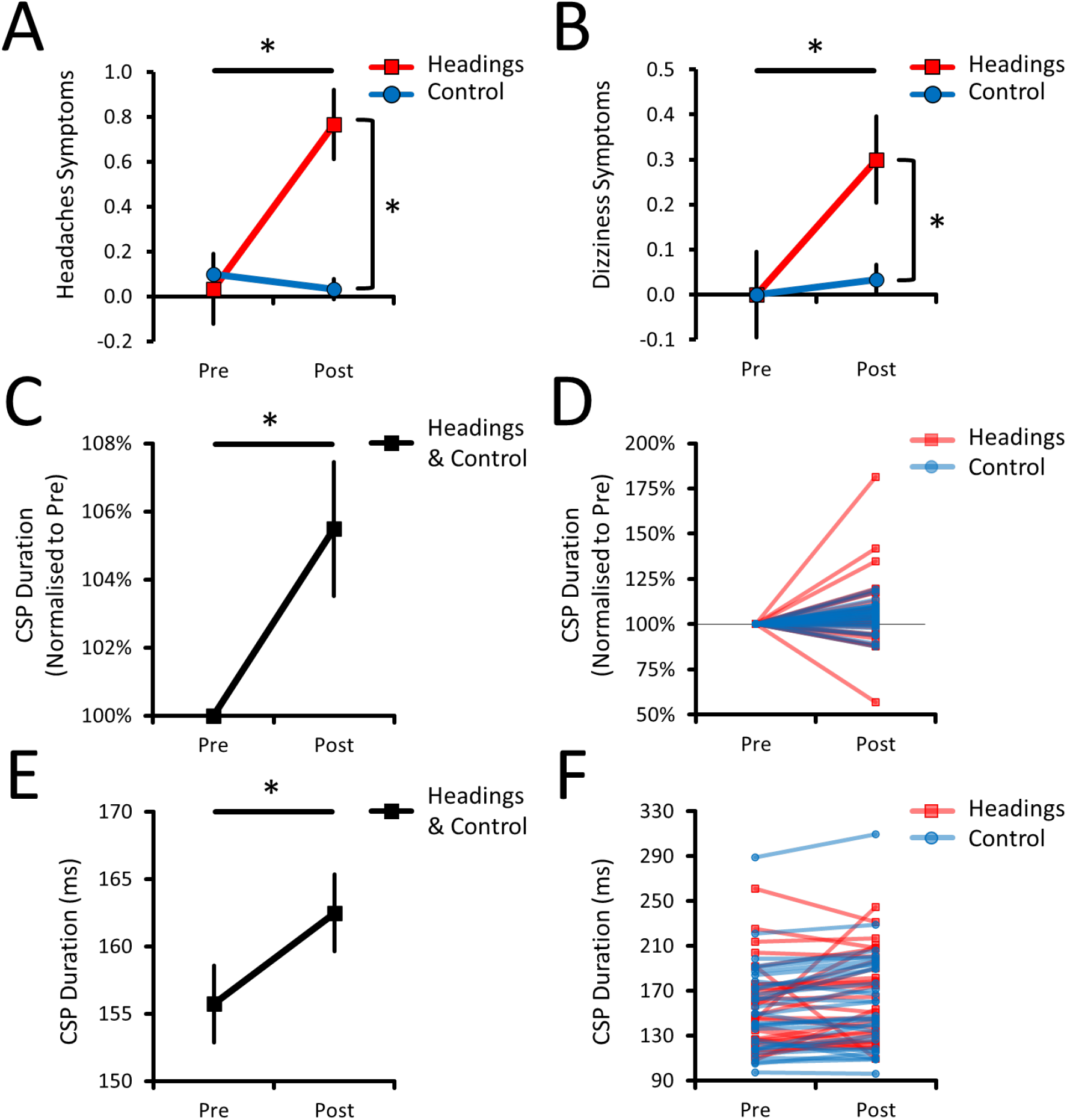
Headaches, Dizziness, and CSP results. **(A)** Headaches and **(B)** Dizziness symptoms, as evaluated with the Rivermead Questionnaire. The Headings group showed greater headaches and dizziness symptoms after headings were performed as compared to the Control group. This indicates that head impacts were experienced in the Headings, but not in the Control group. **(C)** Mean change and **(D)** individual data of CSP duration, depicted as a percentage change from the Pre Measures. The results revealed an effect of Measures (*p* = 0.0087, 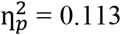), but no interaction between Measures and Groups (*p* = 0.6472, 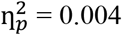), showing that CSP duration lengthened similarly in both groups. **(E)** Mean change and **(D)** individual data of CSP duration, depicted as values in milliseconds. The results also revealed an effect of Measures (*p* = 0.0248, 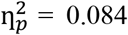) and no interaction between Measures and Groups (*p* = 0.9850, 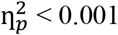), confirming that the results in **(C)** and **(D)** are not a by-product of normalising the CSP duration. For panels (A), (B), (C), and (E), the mean ± 95% CIs are shown. Asterisks (*) denote significant differences.

For dizziness symptoms (Figure 2B), the results also revealed a Measures * Groups interaction (F_(1,58)_ = 6.676, *p* = 0.0123, 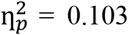). A breakdown of the interaction revealed that dizziness symptoms increased from Pre to Post for the Headings (*p* = 0.0166, Cohen’s dz = 0.561), but not for the Control group (*p* = 1.000, Cohen’s d = 0.183). The Headings and Control groups did not differ at Pre (*p* = 1.000, Cohen’s d = 0.000), but Headings showed greater dizziness symptoms than the Control group at Post (*p* = 0.0122; Cohen’s d = 0.667). This result confirms that participants experienced dizziness symptoms when heading footballs but not in the control group.

Concerning the remaining symptoms (Table 1), the results selectively revealed an effect of Groups (F_(1,58)_ = 4.372, *p* = 0.0409, 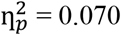) and a marginal Measures * Groups interaction (F_(1,58)_ = 2.866, *p* = 0.0959, 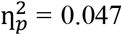) for “Taking longer to think” symptoms. Whilst the effect of Groups revealed that participants in the Headings group “took longer to think” than in the Control group (p = 0.0409, Cohen’s d = 0.540), the marginal Measures * Groups suggests that this difference was apparent at Post (*p* = 0.0430, Cohen’s d = 0.584) but not at Pre (*p* = 0.5702, Cohen’s d = 0.151). Analyses of all the other symptoms revealed no effect of Measures (all F_(1,58)_ < 2.000, all *p* > 0.1626, all 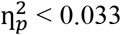), no effect of Groups (all F_(1,58)_ < 1.851, all *p* > 0.1789, all 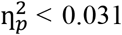), and no Measures * Groups interaction (all F_(1,58)_ < 2.610, all *p* > 0.1116, all 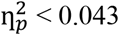). Overall, these results suggest that heading footballs made participants “take longer to think” as compared to the control group. Furthermore, they confirm the absence of meaningful differences within and between groups in the remaining items of the Rivermead Questionnaire.

### CSP duration lengthened in both the Headings and Control groups

A 2 Measures (Pre, Post) * 2 Contraction Levels (80% MVC, 40% MVC) * 2 Groups (Headings, Control) mixed ANOVA was conducted to analyse normalised CSP duration (Figure 2C and D). The results revealed a main effect of Measures (F_(1,58)_ = 7.363, *p* = 0.0087, 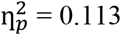), but no Measures * Groups interaction (F_(1,58)_ = 0.212, *p* = 0.6472, 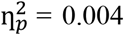). This shows that CSP duration similarly lengthened from Pre to Post for both the Headings (6.4 ± 3.7%) and Control groups (4.6 ± 1.3%). Also, there was no effect of Contraction Levels (F_(1,58)_ = 2.527, *p* = 0.1173, 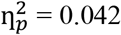), no effect of Groups (F_(1,58)_ = 0.212, *p* = 0.6472, 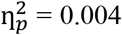), no Measures * Contraction Levels interaction (F_(1,58)_ = 2.527, *p* = 0.1173, 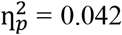), no Contraction Levels * Groups interaction (F_(1,58)_ = 0.231, *p* = 0.6325, 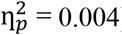), and no three-way interaction (F_(1,58)_ = 0.231, *p* = 0.6325, 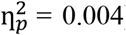). Overall, this analysis shows that heading footballs increased CSP duration, but not more so than in a control group that did not head the ball. It also shows that CSP duration did not differ between the different Contraction Levels.

To confirm that these results were not a by-product of normalising the CSP duration data, the same mixed ANOVA was conducted on the non-normalised CSP duration data (in ms; Figure 2E and F). The results also revealed a main effect of Measures (F_(1,58)_ = 5.308, *p* = 0.0248, 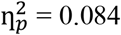) and no Measures * Groups interaction (F_(1,58)_ < 0.001, *p* = 0.9850, 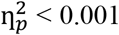). This analysis shows that CSP duration similarly lengthened for both groups from Pre to Post (Headings: 6.8 ± 10.8ms; Control: 6.7 ± 3.9ms). Also, both groups did not differ at Pre (Headings: 157.3 ± 13.1ms; Control: 154.1 ± 14.7ms; *p* = 0.6865, Cohen’s d = 0.082) or Post (Headings: 164.1 ± 13.7ms; Control 160.8 ± 16.2ms; *p* = 0.7576, Cohen’s d = 0.079), suggesting that both groups had homogenous CSP duration. The results also revealed no effect of Contraction Levels (F_(1,58)_ = 0.009, *p* = 0.9245, 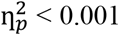), no effect of Groups (F_(1,58)_ = 0.104, *p* = 0.7477, 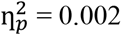), no Measures * Contraction Levels (F_(1,58)_ = 2.671, *p* = 0.1076, 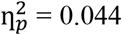), no Contraction Levels * Groups interaction (F_(1,58)_ = 1.626, *p* = 0.2073, 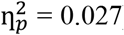), and no three-way interaction (F_(1,58)_ = 0.004, *p* = 0.9509, 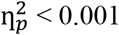), confirming that the above results are not a by-product of normalising the CSP duration data.

## Discussion

This work’s objective was to test the hypothesis that CSP would lengthen in response to football headings (as in Di Virgilio et al., 2016) *and* as compared to a control group. However, the results do not support this hypothesis. Namely, the results first revealed increased headaches and dizziness symptoms in the Headings compared to the Control group, indicating that sub-concussive head impacts were experienced in the Headings group only. The results then revealed that CSP duration similarly lengthened in both the Headings and Control groups, suggesting that head impacts were not the cause of the observed increase in CSP duration. One implication is that changes in brain excitability – as assessed by CSP duration – are not a reliable indicator of acute sub-concussive brain injury.

### Increased headaches and dizziness symptoms after heading footballs

One novel result of this work is that performing football headings (Headings group) increased subjective headaches and dizziness symptoms as compared to hitting the ball with the dominant lower limb (Control group), indicating that head impacts were experienced in the Headings group only. Interestingly, the symptom “taking longer to think” was also marginally increased following the head impacts as compared to control, aligning with previous work suggesting that football headings deteriorate cognitive performance (Di Virgilio et al., 2016; Dioso et al., 2022). The present results extend previous work (Di Virgilio et al., 2016, 2019; McNabb et al., 2020) by showing that sub-concussive head impacts also induce subjective effects that can be monitored using the Rivermead Questionnaire. Overall, these results indicate that head impacts were experienced in the Headings group only and suggest that – in absence of a validated questionnaire developed for that purpose – the Rivermead Questionnaire can be used to assess some of the subjective effects of sub-concussive head impacts.

### CSP duration similarly lengthened for both groups

The main result of this work is that CSP duration lengthened similarly for both the Headings and Control groups, suggesting that acute head impacts are not the cause of the CSP lengthening. Moreover, CSP duration was not affected by the strength of the muscle contraction performed by the participant (80% MVC vs 40% MVC), aligning with previous work showing that different levels of voluntary muscle contraction do not affect CSP durations (Kojima et al., 2013). Whilst the present results replicated the CSP lengthening reported by Di Virgilio et al. (2016), the lack of difference between the Headings and Control groups indicates that such CSP lengthening cannot be attributed to performing football headings *per se*. Therefore, a factor unrelated to head impacts – but present in both the Headings and Control groups – must account for the present CSP lengthening or obscure differences between the groups.

The Control group was designed to match the exercise levels – albeit low in absolute terms – of the Headings group. Thus, one possibility is that the performing comparable exercise levels increased CSP duration similarly for both groups. High and low-intensity aerobic exercise alone can reduce intracortical inhibition in M1 (Stavrinos & Coxon, 2016; Yamazaki et al., 2019) but can also increase GABA concentrations in cortical motor areas (Coxon et al., 2018). Although this evidence makes it unclear if exercise should shorten or lengthen CSP duration, it nonetheless suggests that groups performing similar levels of exercise will show similar changes in CSP duration. In opposition, Di Virgilio et al. (2019) reported that three 3-min sparring bouts increased CSP duration as compared to a mock-sparring control group, suggesting that head impacts should increase CSP duration even if exercise levels are matched. However, Di Virgilio et al. (2016) did not control for exercise levels, making it unclear if the increased CSP duration they reported was due to heading footballs or to the exercise levels achieved during the intervention. Nonetheless, one obstacle for future studies will be to isolate the effects of exercise from those of sub-concussive head impacts, as the two usually co-occur and can therefore confound each other (Tremblay et al., 2018). Overall, one possibility is that raising the exercise level above the resting (sedentary) state CSP are typically recorded in – alone – accounts for the similar CSP lengthening in both the Headings and Control groups.

Another possibility is that changes in CSP duration between different groups can only be observed – or enhanced – when groups have different history of concussions (for a review, see Scott et al., 2020). For instance, De Beaumont et al. (2007) assessed CSP duration at rest in varsity athletes with a concussion history (between 2 to 5 concussions, which occurred more than 9 months before testing) and in control participants with no concussion history. Their results showed greater CSP duration in the varsity athletes as compared to the control participants, suggesting that having a concussion history could predispose to CSP duration lengthening in response to sub-concussive head impacts (see McNabb et al., 2020). De Beaumont et al. (2007) also reported that concussion severity positively correlated with CSP duration whereas the number of experienced concussions did not, suggesting that a history of severe concussions is an important confounding factor. Here, the participants all reported having no history of concussion in the 5 years before they participated in this study, which was shown to be sufficient to restore levels of inhibition and excitation in M1 (Tremblay et al., 2014). As a result, one possibility is that the present lack of difference in CSP duration stems from a similar lack of concussion history between the Headings and Control groups.

## Conclusion

This work shows that heading footballs did increase CSP duration, but not more so than in a control group that did not head footballs, suggesting that heading footballs did not account for the acute increases in corticomotor inhibition. By the same token, this also suggests that changes in CSP duration do not reliably reflect acute brain injuries, as confounding factors other than head impacts can cause CSP to lengthen. Future studies should carefully control factors such as the levels of exercise performed and the history (and severity) of concussion over (at least) the last 3 years (Tremblay et al., 2014). Importantly, the present methods can be used as an experimental model to evaluate the effects of sub-concussive head impacts on brain functions. Future studies could use the present design to collect data in even larger cohorts (n > 100) and could include wet (inflammatory) biomarkers to assess additional dimensions of brain injury in response to sub-concussive head impacts (see Sandmo et al., 2022).

## Acknowledgments

R.H. was funded with a scholarship from the *Fonds de Recherche du Québec – Nature et Technologie* (Québec, Canada).

